# Microfluidic On-demand Engineering of Exosomes towards Cancer Immunotherapy

**DOI:** 10.1101/478875

**Authors:** Zheng Zhao, Jodi McGill, Mei He

## Abstract

Extracellular Vesicles (EVs), particularly exosomes (30-150 nm), are an emerging delivery system in mediating cellular communications, which have been observed for priming immune responses by presenting parent cell signaling proteins or tumor antigens to immune cells. Therefore, preparation of antigenic exosomes that can play therapeutic roles, particularly in cancer immunotherapy, is emerging. However, standard benchtop methods (e.g., ultracentrifugation and filtration) lack the ability to purify antigenic exosomes specifically among other microvesicle subtypes, due to the non-selective and time-consuming (>10 h) isolation protocols. Exosome engineering approaches, such as the transfection of parent cells, also suffer from poor yield, low purity, and time-consuming operations. In this paper, we introduce a streamlined microfluidic cell culture platform for integration of harvesting, antigenic modification, and photo-release of surface engineered exosomes in one workflow, which enables the production of intact, MHC peptide surface engineered exosomes for cytolysis activation. The PDMS microfluidic cell culture chip is simply cast from a 3D-printed mold. The proof-of-concept study demonstrated the enhanced ability of harvested exosomes in antigen presentation and T cell activation, by decorating melanoma tumor peptides on the exosome surface (e.g., gp-100, MART-1, MAGE-A3). Such surface engineered antigenic exosomes were harvested in real-time from the on-chip culture of leukocytes isolated from human blood, leading to much faster cellular uptake. The activation of gp100-specific CD8 T cells which were purified from the spleen of 2 Pmel1 transgenic mice was evaluated using surface engineered exosomes prepared from muring antigen presenting cells. Antigen-specific CD8 T cell proliferation was significantly induced by the engineered exosomes compared to native, non-engineered exosomes. This microfluidic platform serves as an automated and highly integrated cell culture device for rapid, and real-time production of therapeutic exosomes that could advance cancer immunotherapy.

## Introduction

Extracellular Vesicles (EVs), especially exosomes in the nano-size range of 30 ∼150 nm, have shown important roles in intercellular communications in recent decades^1–4^. Immune cell-derived exosomes have been well documented in the regulation of immune stimulation or suppression^5–6^, driving inflammatory^7^, autoimmune^8^ and infectious disease pathology^9–11^. Due to the formation of exosomes beginning with the creation of endosomes as the multivesicular bodies (MVBs), exosomes differ from other cellular membrane-derived microvesicles^12^ in terms of molecular contents. Therefore, exosomes contain specific proteins and nucleic acids and represent their parent cell status and functions at the time of formation ^13–14^. Among many subtypes of exosomes, immunogenic exosomes with an intrinsic payload of MHC class I and II molecules and other co-stimulatory molecules are able to mediate immune responses^15–16^, which opens up opportunities for the development of novel cancer vaccines and delivery in immunotherapy^17–26^.

Compared to other nano-sized delivery systems, such as lipid, polymers, gold, and silica material^27–32^, exosomes are living-cell derived, highly biocompatible nano-carriers with intrinsic payload, and exhibit much stronger flexibility in loading desired antigens for effective delivery^33^. Exosomes also eliminate allergenic responses without concerns of carrying virulent factors and avoid degradation or loss during delivery^34–35^. However, the development of exosome-based vaccines is hindered by substantial technical difficulties in obtaining pure immunogenic exosomes^36^. The diverse subtypes of exosomes could confound the investigation on differentiating different cellular messages. On the other hand, molecular engineering of exosomes through either membrane surface or internal loading could provide an untapped source for developing novel antigenic exosomes^16^. Bioengineered exosomes as emerging immunotherapeutics have gained substantial attention in developing a new generation of cancer vaccines, including recent phase-II trial using IFN-DC-derived exosomes loaded with MHC I/II restricted cancer antigens to promote T cell and natural killer (NK) cell-based immune responses in non-small cell lung cancer patients^21, 37–41^. Unfortunately, current exosome engineering approaches, such as the transfection or extrusion of parent cells and membrane permeabilization of secreted exosomes, suffer from poor yield, low purity, and time-consuming operations^38, 42–45^. Therefore, in this paper, we introduce a facile, 3D-printing molded PDMS microfluidic culture chip for solving this bottleneck problem. Due to the intrinsic features in automation and high-efficient mass transport, microfluidic systems overcome many of the drawbacks of benchtop systems and show superior performance in isolating^46–51^, detecting and molecular profiling exosomes^52–56^. However, the potential of the microfluidic platform for molecular engineering of exosomes has not been well explored yet^38^. Herein, we developed the 3D molded PDMS microfluidic device which enables real-time harvesting, antigenic modification, and subsequent photo-release of intact, engineered antigenic exosomes on-demand. Specifically, we introduced a novel magnetic-nanoparticles functionalized with photo-cleavable, peptide affinity probe for capturing and on-demand releasing MHC-I positive exosomes via a light trigger. After antigenic loading of MHC-I binding peptides on the surface of captured exosomes, the photo-release of modified exosomes in the microfluidic system can be well controlled spatially and temporally with ∼95% release efficiency. Presently, the reported work on processing exosomes via microfluidic approaches are either in small quality or bound to solid surface/particles, and unable to release without extra capture probes/tags or stay intact for downstream therapeutic preparations^57–58^. To the best of our knowledge, no comparable work has been reported for engineering cell-secreted antigenic exosomes in real time for advancing cancer immunotherapy. Such a functional-streamlined microfluidic cell culture system allows antigenic engineering of exosomes either through mediating their parent cell growth using stimulations, or direct molecular engineering on the surface of produced exosomes. We studied several tumor antigenic peptides (e.g., gp-100, MAGE-A3, and MART-1) which are commonly used in developing cancer vaccines but difficult in delivery due to degradation. Our microfluidic system showed high-efficiency in engineering immunogenic exosomes (MHC I+), meanwhile, photo-releasing the intact functional exosomes downstream. We tested the cellular uptake of engineered exosomes by using antigen presentation cells, which displayed much-improved internalization ability compared to native, non-engineered exosomes. We also assessed the immunogenicity of engineered exosomes for activating transgenic mouse isolated antigen-specific CD8 T cells, demonstrating their capacity to induce peptide-specific T cell proliferation and prove the viability and functionality of engineered exosomes towards application in cancer immunotherapy. This facile and low-cost microfluidic platform not only provides a novel enabling strategy for high-efficient production of purified, enriched therapeutic exosomes, but also serves as an investigation tool for understanding roles of variable peptide-engineered exosomes in antitumor immune responses and cancer immunotherapy (See Fig. 1).

**Fig. 1.**
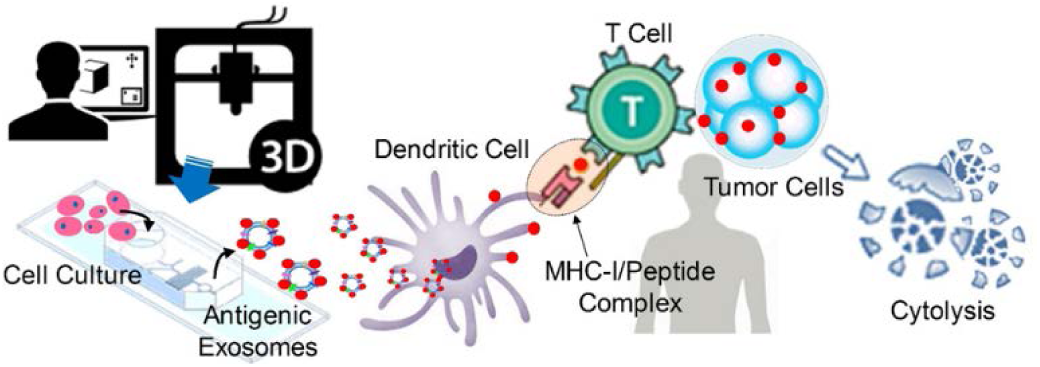
Illustration of 3D-printing molded PDMS microfluidic culture chip for streamlined engineering of antigenic exosomes employed in activating anti-tumor responses.

## Experimental

### 3D Printing and Fabrication of Microfluidic Culture Chip

Three pieces of the molds were used for microfluidic PDMS chip fabrication, including a base, the side-wall molding, and top magnet holder. Molds were designed by using the SolidWorks^®^ 2017 and printed out by the 3D printer (Project 1200 from 3D Systems). The finest structures printed by the 3D printer is in 30 μm. The height of the microfluidic channel is molded at 50 μm. The cylindrical cell culture chamber is molded in 1000-μm diameter and 500-μm height. All 3D-printed molds were sputtering coated with palladium in a thickness of 20 nm for easy release of molded-PDMS. Three pieces of molds were assembled to cast the PDMS microfluidic cell culture device as shown in Figure S1, which was cast by a 10:1 ratio of Dow SYLGARD^TM^ 184 silicone solution (Sigma-Aldrich) and incubated at the temperature of 40 °C for 6 hours. After a complete cure, the molded PDMS polymer can be peeled off easily. The molded cell culture chamber is open to the air and allows a PDMS-made plug to cap on the top. Chip inlets and outlets were formed by punching holes in 0.75 mm diameter. Piranha solution-cleaned glass slides and the PDMS layer were both treated with high-voltage plasma for at least 30 seconds, and then bond on the hot plate at the temperature of 40 °C for 5 mins. The formed microfluidic cell culture chip was cleaned by DI water, and then sterilized using autoclave at 121 °C for 30 mins.

### On-chip Cell Culture and Exosome Collection, Engineering, and Releasing

The glass coverslip (8 mm) was autoclaved at 121 °C for 30 mins and treated with 500 uL of 0.1 mg/mL Poly-D-Lysine Hydrobromide (MP Biomedicals) before putting into the 24-well plate for cell seeding. Murine JAWSII cells (ATCC) or leukocytes isolated from human blood buffy coats (Innovative Research) were seeded in the 24-well plate which contains coverslips at the bottom of wells in the biosafety cabinet.

For surface engineering of cell-secreted exosomes, a solution of β2-microglobulin at 20 μm/mL (Sigma-Aldrich, either human or murine) with synthesized tumor antigenic peptides at 100 μm/mL (MAGE-A3 from Genscript inc.; gp-100 and MART-1 were synthesized by KU Molecular Probe Core, see supplemental information for synthesis conditions, quality characterization and sequences) were prepared in the 1× PBS buffer as the antigenic loading buffer. The concentration of β2-microglobulin was experimentally optimized based on the reference^16^. As shown in Fig. 2, by closing the B-inlet, antigenic loading buffer can be pumped through the A-inlet and the washing buffer can be pumped through the C-inlet into microfluidic channels at the volume flow rate of 1 μL/min or 0.1 μL/min for 10 mins. The washing step can be performed from both A-inlet and C-inlet at the volume flow rate of 1 μL/min for 15 mins. By taking off the magnet from the underneath of collection microchamber, the UV light via a UV objective can be projected to the magnetic beads aggregation area for photo-release of captured exosomes. The streptavidin magnetic beads (500 nm, Ocean Nanotech Inc) were conjugated with the Biotin-PC-NHS photo-cleavable linker (BroadPharm Inc.) and peptide probes per vendor’s instruction. By applying another washing step from both A-inlet and C-inlet at the volume flow rate of 1 μL/min for 20 mins, the photo-released exosomes can be collected at the outlet of a microfluidic chip for subsequent in vitro or in vivo study.

**Fig. 2.**
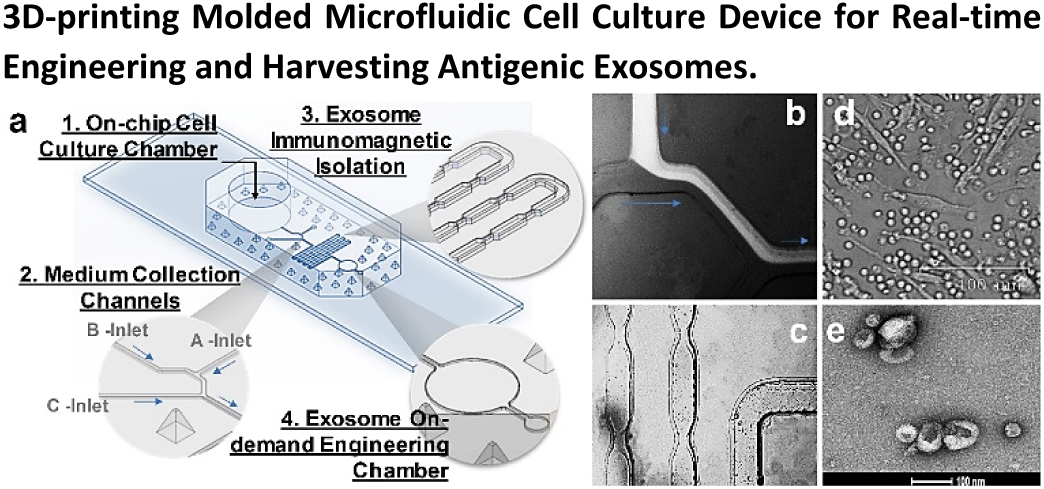
a) Illustration of 3D-printing molded microfluidic culture device for engineering immunogenic exosomes directly from on-chip cultured cells in real time. b) Fluorescence dye solution for showing the flow of immunomagnetic beads (A-Inlet) mixing with cell culture media (B-Inlet) eluted from cell culture chamber. c) Bright-field microscopic image showing the immunomagnetic microbeads mixing process for isolating exosomes in the serpentine microchannel. d) Bright-field microscopic image showing the morphology of on-chip cultured leukocytes. e) SEM image showing the engineered exosomes released out of the chip.

### Exosomes Staining and Cellular Uptake

The microfluidically collected 20 μL exosomes were added to the ultracentrifuge tube and diluted to the final volume of 1 mL for centrifugation (Thermo Scientific™ Sorvall™ MTX) under 1,500 rcf for 30 mins. The supernatant was transferred to a fresh ultracentrifuge tube, and then centrifugated at the speed of 100,000 rcf for 1 hour. The cleaned exosomes were stained by using the PKH67 Green Fluorescent Cell Linker Midi Kit (Sigma-Aldrich) per vendor’s instruction. For removing free dyes after exosome membrane dye labeling, 2 mL of FBS (exosome depleted) medium was added to quenching the free dye. The 1.5 mL of 0.971 M sucrose solution was prepared in complete media with total volume of 10 mL for density gradient centrifugation at 100,000 rcf for 1 hour. The supernatant was discarded after washing off the free dye. The 2 mL of 1× PBS was used to re-suspend the pellet. The 1 μL of Penicillin-Streptomycin (ATCC) was used to sterilize collected exosomes in solution. The collected final exosomes can be stored at 4°C for less than 1 week, and at −20°C for up to one month.

For cellular uptake experiments, the human THP-1 cell line (ATCC) was cultured using ATCC-formulated RPMI-1640 Medium plus 10 % exosome-depleted FBS. The cells used for uptaking exosomes were cultured at the density of 5×10^5^/mL. The 20 μL native, non-engineered exosomes (NE) were added into 5 wells as the control group. The other 20 μL surface engineered exosomes (EE) were added into 5 wells separately as the experimental group. One extra well was set as the negative control. The incubation time intervals were set at 0 hour, 0.5 hour, 1 hour, 2 hours, 3 hours, and 4 hours. At each time point, cells were collected and suspended into 200 μL media for the cytocentrifugation at the speed of 400 rpm for 4 mins. The Fixative Solution (ThermoFisher) was used to cytospin cells and incubated at room temperature for 18 mins. The 100 μL of 1× PBS buffer was used to wash the fixed cells for three times. The slide was kept air-dry without any remaining water drops. The DAPI (ThermoFisher) was used to stain the cell nucleus per vendor’s instructions. The ProLong™ Gold Antifade Mountant (ThermoFisher) was used and then covered with the coverslip without trapping any air bubbles. The prepared slides can be stored at room temperature for 24 hours before microscopic imaging.

For IFN-ɤ stimulation experiment, the gp-100 engineered exosomes and native, non-engineered exosomes were incubated with leukocytes in 96-well plate for monitoring stimulated cytokine secretion due to the antigen-specific immuno-responses. The red blood cell lysis, and human leukocytes culture and stabilization follow the standard protocols elsewhere (Innovative Research). The four different stimulation level of harvested exosomes were used (0 μL, 5 μL, 10 μL, 20 μL), and 20 μL was used as the best effective dose. The IFN-ɤ concentration from each condition (24 hr, 36 hr, and 48 hr) was measured using the IFN gamma ELISA Kit from Thermo Fisher with pokeweed mitogen protein as the positive stimulation control, per vendor’s instruction. Three repeats were measured for calculating RSD.

### In vitro CD8+ T Cell Activation and Flow Cytometry Analysis

Female, CD8 T cell transgenic mice, aged 6-7 weeks, were purchased from Jackson Laboratories (B6.Cg-Thy1a/Cy Tg(TcraTcrb)8Rest/J). CD8 T cells from this strain express a transgenic T cell receptor that is specific for the gp-100 peptide of the premelanosome 17 protein. The murine monocyte JAWS-II cell line was purchased from ATCC. JAWS cells were cultured per ATCC recommendation and were maintained in alpha-MEM with ribonucleosides, deoxyribonucleosides, 4 mM L-glutamine, 1 mM sodium pyruvate, 5 ng/ml murine GM-CSF and 20% fetal bovine serum. For T cell co-culture studies, JAWS cells and T cells were maintained in complete RPMI composed of RPMI-1640 (Gibco, Carlsbad, CA) supplemented with 2 mm l-glutamine, 25 mm HEPES buffer, 1% antibiotic– antimycotic solution, 50 mg/mL gentamicin sulphate, 1% non-essential amino acids, 2% essential amino acids, 1% sodium pyruvate, 50 μm 2-mercaptoethanol and 10% (volume/volume) fetal bovine serum.

For T cell co-culture experiments, JAWS cells were activated 48 hours prior to the start of the experiment. JAWS cells were seeded at a concentration of 4×10^4^ cells/well in cRPMI in a 96-well plate and were stimulated overnight at a concentration of 200 ng/mL Lipopolysaccharide (LPS). Parallel wells of JAWS cells remained unstimulated and were incubated overnight in media only. JAWSII cells were then washed twice with warm cRPMI.

Transgenic CD8 T cells were purified by magnetic cell separation per manufacturer’s instructions (Miltenyi Biotech) and then labeled with Cell Trace Violet (Life Technologies). CD8 T cells were enumerated and plated at a concentration of 2×10^5^ cells/well in cRPMI in wells alone (T cells only), with activated JAWSII cells or with normal JAWSII cells. Surface engineered exosomes were then added to the CD8 T cell cultures at increasing ratios of exosomes: dendritic cells (25, 50 and 100). Negative control wells did not receive exosomes. Positive control wells were stimulated with 5 ug/mL Conconavalin A. The cells and exosomes were co-cultured for 5 days. Cell cultures were then labeled with anti-mouse CD3 monoclonal antibody and anti-mouse CD8 monoclonal antibody (both from BD Biosciences) and CD8 T cells were analyzed by flow cytometry for Cell Trace Violet dilution. Cells were collected by a BD FACS Canto flow cytometer and the data were analyzed using FlowJo v10 (TreeStar).

## Results and Discussions

### 3D-printing Molded Microfluidic Cell Culture Device for Real-time Engineering and Harvesting Antigenic Exosomes

In contrast to microfabrication conducted in a clean room, we introduce a facile and low-cost approach for making a PDMS microfluidic cell culture device via a 3D-printed mold (see Figure s1) with good molding reproducibility (RSD ∼3%) and precision. This microfluidic culture device contains a cell culture chamber for on-chip growing cells and collecting exosomes in real time from eluted culture medium at downstream. The cell culture chamber is open to air on the top for applying a PDMS-made, finger-push plug, which can produce pressure for flowing medium to downstream collection microchannel, as well as the medium exchange. The collection microchannel (B-Inlet) interconnects culture chamber at the bottom and an A-Inlet (200-μm wide and 200-μm high) for introducing immunomagnetic isolation beads to mix with eluted culture medium, as shown in Fig. 2. The consecutive narrow-neck microstructures (250 μm: 75μm ratio) in serpentine microchannel are shown in Fig. 2 exploded isolation area for enhancing the mixing process. The C-Inlet is used to introduce the washing buffer driven by a syringe pump. Fig. 2b demonstrated the mixing process through the A-Inlet and B-Inlet, and then exited to the exosome isolation serpentine channel by imaging a fluorescence dye solution using the fluorescence microscope. Fig. 2c records the immunomagnetic beads mixing process within the serpentine channel. Human blood-derived leukocytes were cultured in the microfluidic culture chamber with the morphology shown in Fig. 2d. Few red blood cells were observed as a cup shape, and the majority of cells were lymphocytes. The harvested exosomes from downstream of the microfluidic cell culture device were characterized by SEM imaging shown in Fig. 2e, after on-chip immunomagnetic capture, isolation, and photo-release. The typical round cup shape in 100 nm was observed from harvested exosomes, which demonstrated the effective collection of cell secreted antigenic exosomes.

### On-demand Photo-release of Surface Engineered Exosomes

The current existing bead-based exosome isolation approach always delivers particles or solid surface-bound exosomes which inhibits further delivery of exosomes for cellular uptake or therapeutic applications. Therefore, we developed the conjugation of a photo-cleavable linker which contains bi-functional groups of biotin and NHS chemistry for anchoring to the streptavidin immunomagnetic bead surface, meanwhile, keeping the other end of NHS group covalently bond with the MHC-I binding peptide via the primary amine, as shown in Fig. 3a. The MHC class I molecules are heterodimers that consist of two polypeptide chain α, one noncovalent interaction domain of β2-microglobulin (β2m), and one α3 domain. The two domains of α1 and α2 are folded to make up a groove for binding to 8-10 amino acid peptides (MHC-I binding peptide). The formed MHC-I/peptide binding complex can be displayed to cytotoxic T cells consequently for triggering an immediate response from the immune system. Once the MHC-I positive exosomes are captured by tumor targeting antigenic (TTA) peptide and retained by immunomagnetic beads within the capture chamber under the magnetic field, the antigenic loading buffer with saturated TTA peptides can be introduced via C-Inlet to completely bind and replace the rest available MHC-I peptide binding sites. This antigenic surface engineering process can substantially enhance the loading amount of TTA peptides to the captured MHC-I positive exosomes and boost surface antigen presentation to activate T-cells.

**Fig. 3.**
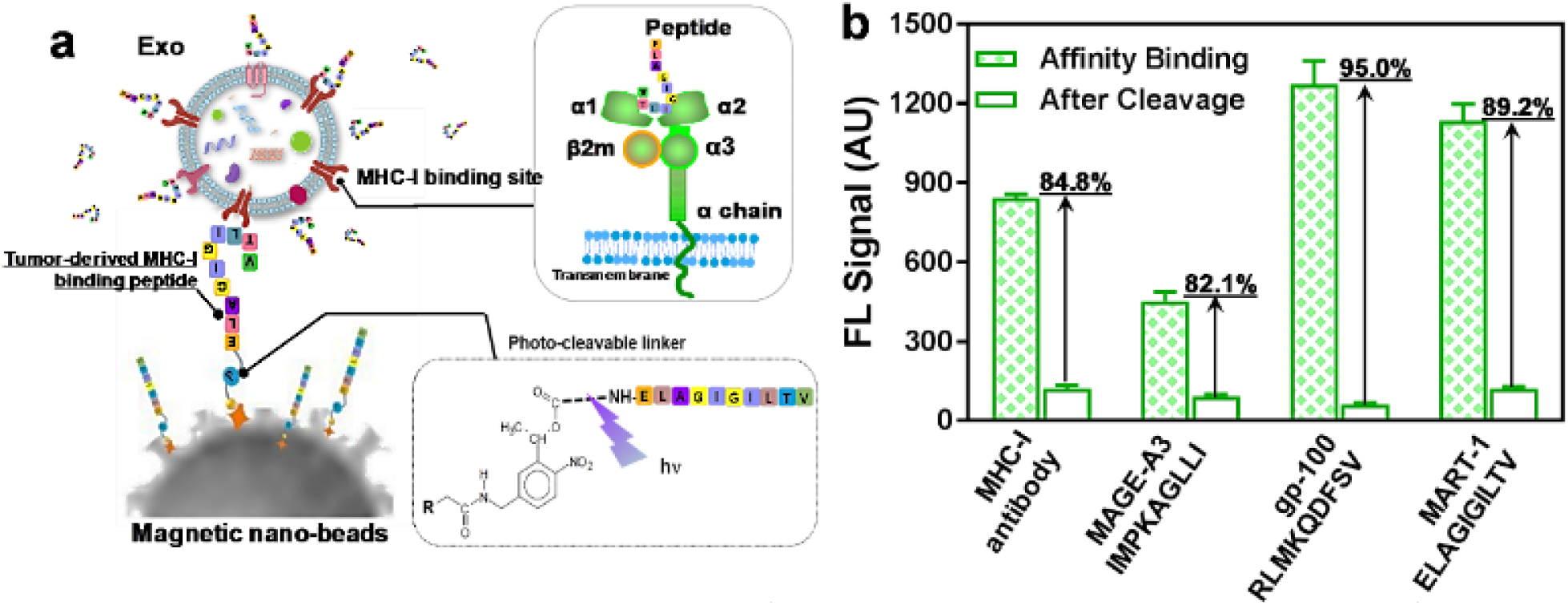
a) The illustration of immunomagnetic capture and on-demand photo-release of MHC-I positive, antigenic exosomes. b) Characterization of three tumor-targeting peptide antigens conjugated with photo-cleavable immunomagnetic beads for binding and photo-release of fluorescence-labeled immunogenic exosomes. The MHC-I antibody is used as the positive control. The error bar shows the three repeats with average measurement (RSD < ∼5%)

For evaluating the binding strength between the MHC-I/peptide complex, several TTA MHC binding peptides were used as the affinity probe for capturing via photo-cleavable linker conjugated immunomagnetic beads, as shown in Fig. 3b. Herein, the MHC-I positive exosomes labeled with fluorescence can be captured via the MHC-peptide affinity binding. By comparing the fluorescence intensity of captured exosomes before and after photo-release, we can define the recovery rate for light-triggered harvesting. The MHC-I antibody serves as the positive control for capturing fluorescence-labeled exosomes and evaluating capacity that can form the MHC-I/peptide complex. As observed in Fig. 3b, both gp-100 and MART-1 TTA peptides showed much higher fluorescence intensity compared to MHC antibody after affinity capture, which indicates the stronger binding complex formed between exo-MHC-I and gp-100 or MART-1 peptides. The gp-100 exhibits the highest recovery rate of 95% after photo-release. Due to the stronger binding capacity of MHC-I/peptide complex, the surface antigen presentation is more effective which can induce higher potency for activating T cell anti-tumor responses^59–60^. We observed that gp-100 exhibited stronger ability to form MHC-I/peptide complex on exosome surface compared to other MART-1 and MAGE-A3 TTA peptides, and this MHC-I peptide binding capacity is even stronger than MHC-I antibody (95% vs 84.8%).

The performance of on-demand photo-release was characterized in Fig. 4a. With the comparison between positive control and negative control, the evaluation of fluorescence-labeled exosomes on capture and photo-release was performed by measuring fluorescence intensity from beads aggregates under an invert fluorescence microscope. The SEM imaging approach was used to confirm the photo-release process as shown in Fig. 4b and c. By comparing the SEM imaging of beads surface before and after photocleavage, there are no identifiable exosome particles presented on the surface of beads, indicating the good photo-release performance. The UV exposure time was characterized as well for reaching 98% photocleavage rate within 8-minute UV exposure (365 nm, ∼ 2mW/cm^2^). We evaluated the size distribution of photo-released, surface-engineered exosomes for comparing with native exosomes, which showed an appropriate size range of exosomes between 50 nm - 200nm, confirming that surface-engineered exosomes are maintaining a good integrity. The side-effect of UV exposure on exosome molecular contents was investigated in Figure s2, which showed non-detectable changes in terms of exosomal proteins, DNAs, and RNAs under 10-minute UV treatment. This result supports the photo-release of surface-engineered exosomes in good integrity and biological activity for subsequent cellular uptake and immune activation.

**Fig. 4.**
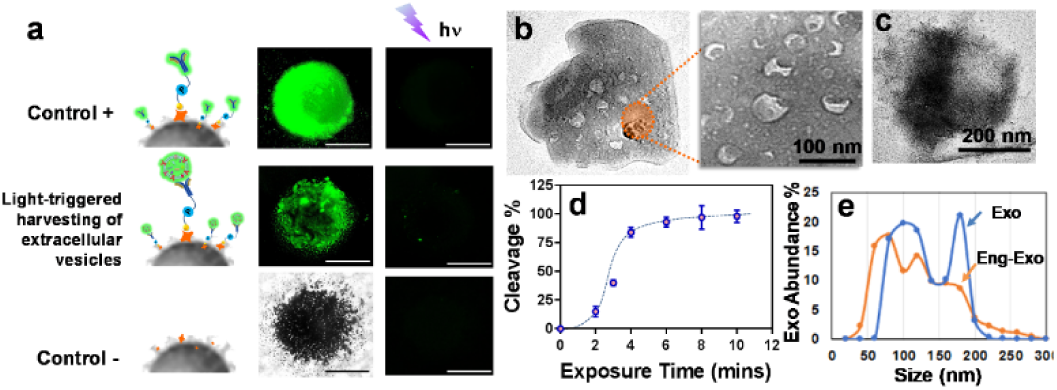
a) Characterization of the performance of on-demand photo-release of captured exosomes from immunomagnetic beads. The positive control is a fluorescence-labeled antibody captured by photo-release immunomagnetic beads. The negative control is the immunomagnetic beads without a photo-cleavable linker. b) The SEM image of the surface of photo-release immunomagnetic beads captured with exosomes. Exosome particles were seen as the cup shape due to the vacuum sample preparation. c) The SEM image of the surface of photo-release immunomagnetic beads after photocleavage. d) Characterization of UV exposure time influence on the photo-cleavage efficiency. The error bar shows the three repeats with average measurement (RSD < ∼5%). e) Nanoparticle tracking analysis of exosome size distribution between photo-released, surface engineered exosomes and native exosomes.

### Immunogenic Potency of Surface Engineered Exosomes

In order to evaluate the potency and integrity of surface-engineered exosomes which are photo-released from the microfluidic cell culture device, we labeled the harvested exosomes with green membrane dye PKH67, and then incubated both gp-100 engineered exosomes and native exosomes with the human monocytic THP1 cells, and monitored cellular uptake at one-hour intervals. After cell fixation and nucleus staining with DAPI, we observed abundantly distributed green dots as exosomes around the cell nucleus (Fig. 5a), which is much more intense than uptake of native exosomes. The cellular uptake begins within one hour for both gp-100 engineered exosomes and native exosomes. However, the uptake capacity and speed is significantly greater for gp-100 engineered exosomes than native exosomes. After 4 hours, both gp-100 engineered exosomes and native exosomes were cleared by the lysosome pathway. This observation indicates that gp-100 engineered exosomes significantly enhanced the cellular uptake from dendritic monocytes by ∼ 2 folds, compared to native exosomes (see Figure s3). We also monitored the expression of Cytokine IFN-ɤ from incubation of gp-100 engineered exosome with human leukocytes using ELISA. Compared with the stimulation from native exosomes, IFN-ɤ expression under gp-100 exosome stimulation is significantly higher by ∼2-fold, with 48-hour continuous monitoring (Fig. 5b). The gray dash line in the Fig. 5b indicates the positive control using pokeweed mitogen protein as the stimulator. The leukocytes morphology upon stimulation was shown in Fig. s4. Compared to the negative control without stimulation (yellow dash line), both pokeweed mitogen and gp-100 engineered exosomes significantly activated the immune lymphocytes and changed the morphology of leukocytes. The gp-100 engineered exosomes induced greater IFN-ɤ production than the control pokeweed mitogen. This result suggests that TTA peptide gp-100 engineered exosomes are rapidly internalized by antigen presenting cells and highly immunogenic for stimulating cytokine production.

**Fig. 5.**
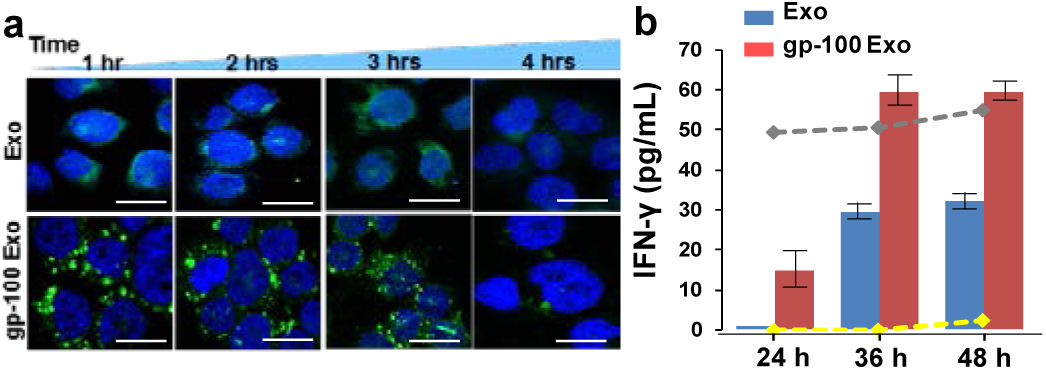
a) The confocal microscopic image of DC uptake of TTA peptide gp-100 surface engineered exosomes, compared with non-engineered native exosomes. The image was taken every one hour for tracking the green fluorescence labeled exosomes uptake by DCs (cell nucleus were stained with DAPI). The scale bar is about 5 μm. b) The release of Cytokine IFN-ɤ from DCs culture under stimulations between native exosomes and gp-100 engineered exosomes. The gray dash line indicates the positive control using pokeweed mitogen protein as the stimulator. The yellow dash line is the negative control without any stimulator. The error bar shows the three repeats with average measurement (RSD is ∼5%).

We further investigated the potency of gp-100 surface-engineered exosomes by testing their capacity to activate antigen-specific CD8+ T cells proliferation. Transgenic, gp100-specific CD8 T cells were purified from the spleen of 2 Pmel1 transgenic mice by magnetic cell sorting and labeled with Cell Trace Violet proliferation dye. The purified T cells were cultured alone (T cells only), mixed at a 3:1 ratio with naïve JAWS cells (an immature dendritic cell line derived from a C57BL/6 mouse), T cells + JAWS cells, or JAWS cells that were activated for 48 hours with 200 ng/mL LPS (T cells + Activated JAWS cells). The surface-engineered exosomes bearing the gp100 peptide were added to the T cell cultures at increasing ratios of exosomes: dendritic cells (25, 50 and 100). The cells and exosomes were co-cultured for 5 days and then CD8 T cells were analyzed by flow cytometry for Cell Trace Violet dilution as a measure of T cell proliferation. As seen in Fig. 6, surface-engineered exosomes alone had no capacity to induce antigen-specific T cell proliferation. However, in the presence of antigen presenting cells, gp100 surface-engineered exosomes stimulated significant CD8 T cell proliferation. The gp-100 exosomes induced the greatest proliferative response when cultured in the presence of LPS-activated JAWS cells, and this response was dose-dependent with ∼30% proliferation rate of CD8+ T cells at the 100:1 ratio of exosomes:dendritic cells. Thus, this result suggests that antigenic surface engineering and photo-release strategy from our developed microfluidic culture device is effective for producing immunogenic exosomes, which could be a powerful tool for developing an effective exosome-based vaccine or delivery agents for priming immunity employed in cancer Immunotherapy.

**Fig. 6.**
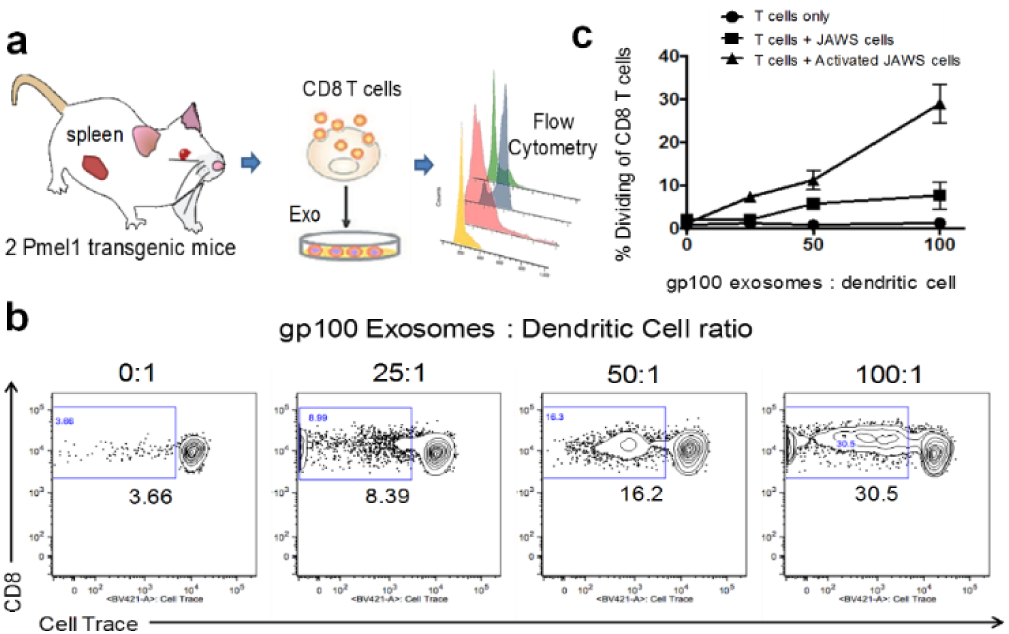
a) Illustration of ex vivo testing of surface-engineered exosomes for activating transgenic mice spleen-derived CD8+ T cells. b) depicts representative flow plots from wells containing T cells + Activated JAWS cells with increasing concentrations of the gp100-engineered exosomes. c) depicts the cumulative data from all three culture conditions showing the CD8+ T cell dividing rate under stimulation. The results are representative of 2 independent experiments with three duplicate wells for each culture condition (RSD < ∼ 5%).

## Conclusions

We demonstrated a simple microfluidic cell culture approach for real-time harvesting, antigenic modification, and photo-release of surface engineered exosomes within one workflow. The microfabrication of the device is straightforward by directly casting PDMS from a 3D printed mold using a consumer-grade 3D printer, without the needs of a clean room. The 3D-molded microstructures contain the z-dimension changes which are very challenging to be made by the conventional photolithographic approach using multi-layer bonding and alignment. The replication process is reproducible with RSD <3% and the variation between designed dimensions is ∼5 μm with 30-μm printing precision.

The integrated antigenic modification and photo-release downstream of cell culture enables real-time investigation of secreted exosomes in a dynamic process. Photo-release of exosomes with enhanced immunogenicity and biological activity has not been reported elsewhere. The recovery rate of released exosomes using our approach is 98% and the exosomes exhibit the strong capacity for uptake by antigen presenting cells, and to induce robust, antigen-specific CD8+ T cell proliferation compared to native exosomes. This developed microfluidic approach can serve as a powerful investigation tool for understanding the roles of variable peptide-engineered exosomes in antitumor immune responses and cancer immunotherapy. Current immunotherapy treatments can only benefit less than 15 % of patients, due to the poorly understood immunity modulation mechanism, and the lack of well-targeted delivery and clinical study designs that are optimized to determine maximum efficacy^61–64^. Immune cell-derived exosomes can be powerful vaccine templates for transferring stimulating factors, antigens, and drugs, which is highly promising by utilizing patient-derived exosomes for developing personalized precision cancer vaccination^40, 65–69^. Our developed microfluidic platform could serve as the powerful technology platform for facilitating this discovery in personalized cancer vaccines.

## Supporting information

## Acknowledgments

The PI Dr. He conceived the research idea, analyzed the data, and drafted the manuscript. The Co-PI Dr. McGill established the animal model, provided the insights into the immunology-related study, and performed testing in Fig 6. Mr. Zhao conducted bench work for Fig. 1 to Fig. 5. We thank the funding support from NIH NIGMS P20GM1000372 and USDA-NIFA 2017-67021-26600. We also thank the KU Microscopy and Analytical Imaging Laboratory for helping with the SEM and TEM imaging, and the KU Molecular Probes Core (KU-MPC) for peptide synthesis.

