## Supplementary material for "Microfluidic On-demand Engineering of Exosomes towards Cancer Immunotherapy"

Zheng Zhao[a], Jodi McGill[b], and Mei He[a,c*]
[a.] Department of Chemical and Petroleum Engineering, Bioengineering Program, University of Kansas, Lawrence, Kansas, USA
[b.] Department of Veterinary Microbiology and Preventive Medicine, Iowa State University, Iowa, USA
[c.] Department of Chemistry, University of Kansas, Lawrence, Kansas, USA

### Supplemental Information

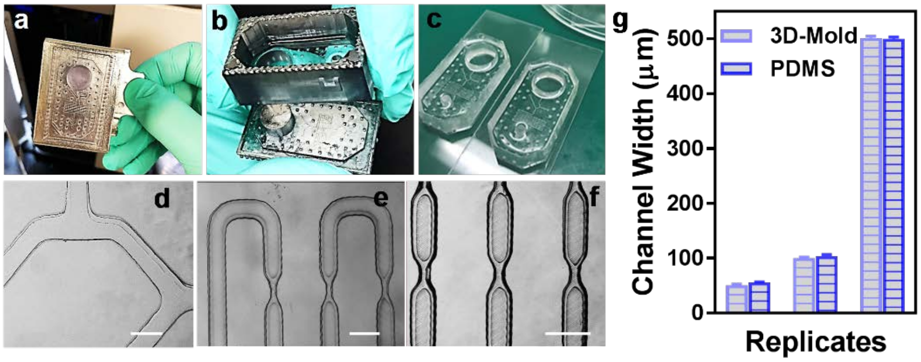


**Figure s1**. Illustration of 3D printing approach for one-step producing 3D mold and replicating PDMS microfluidic device integrated with cell culture and downstream exosome isolation, surface engineering, and on-demand photo release.

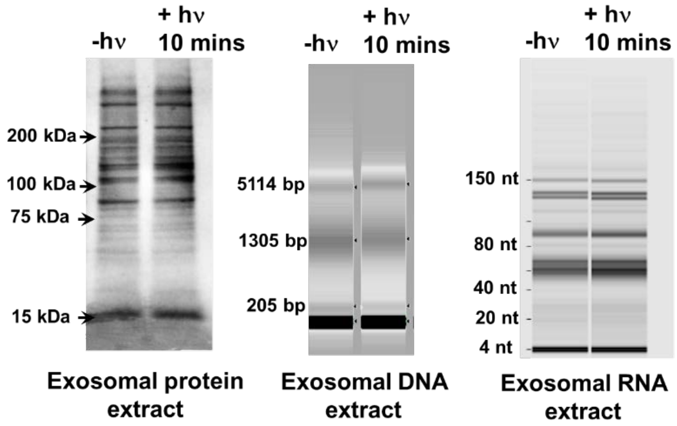


**Figure s2**. Investigation of the side-effect of UV exposure on exosome molecular contents in terms of proteins, DNAs and RNAs.

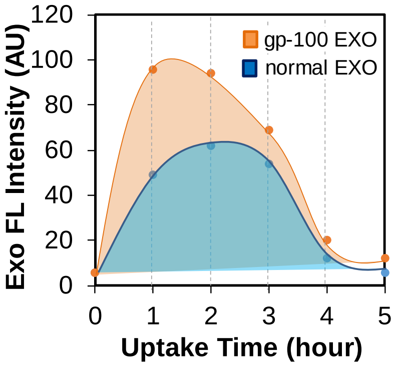


**Figure S3**. The fluorescence intensity analysis for showing the cellular uptake rate of gp-100 engineered exosomes and native exosomes.

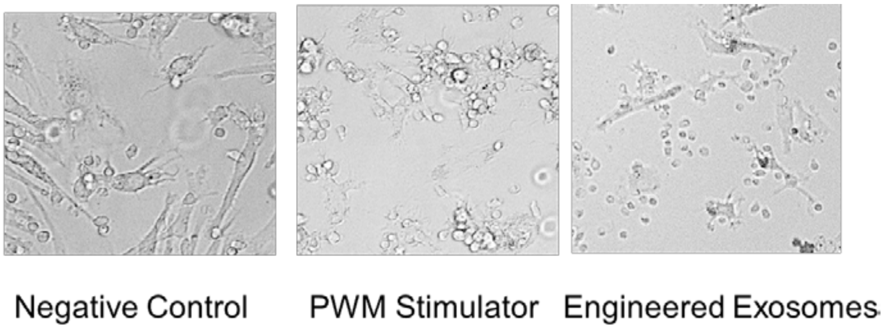


**Figure s4**. Human leukocytes culture under different stimulation conditions: 1) negative control is the leukocytes without any stimulation; 2) PWM protein stimulation as the positive control; 3) The gp-100 engineered exosome stimulation.

**Tumor peptide synthesis and characterization**

The protocols follow standard Fmoc chemistry. The peptides where cleaved using a solution of 92.5:2.5:2.5:2.5 TFA:TIPS:H2O:DODt and the crude peptides where purified using preparative HPLC (gradients of water/ acetonitrile (90:10 to 0:100 containing 0.1% TFA over 40 min) and lyophilized to obtain white powder. Analytical HPLC traces were acquired using an Agilent 1100 quaternary pump and a Hamilton PRP-1 (polystyrene-divinylbenzene) reverse phase analytical column (7 µm particle size, 4 mm x 25 cm) with UV detection at 210 nm. The eluents were heated to 45 °C to reduce separation of rotational isomers, and elution was achieved with gradients of water/ acetonitrile (90:10 to 0:100 containing 0.1% TFA) over 20 min. Low-resolution mass spectra (LRMS) were obtained using a Waters Micromass ZQ 4000 instrument with ESI+ ionization.

**Peptide gp-100 Sequence:** RLMKQDFSV
**Chemical Formula:** C49H82N14O14S1
**Molecular Weight:** 1123.33

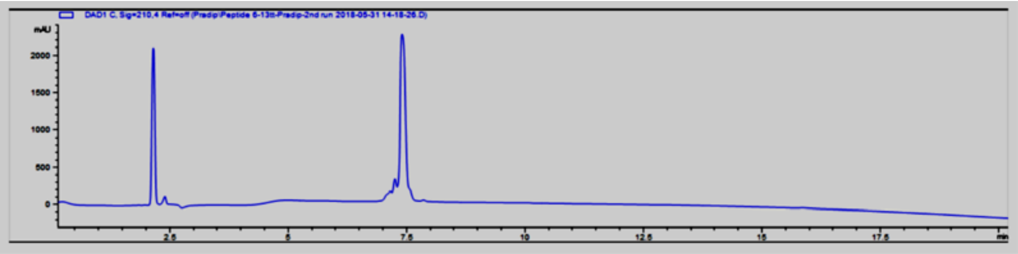

Analytical HPLC profile of sythesized peptide gp-100. Retention time = 7.51min (monitored at 210 nm). Purity > 90% by HPLC.

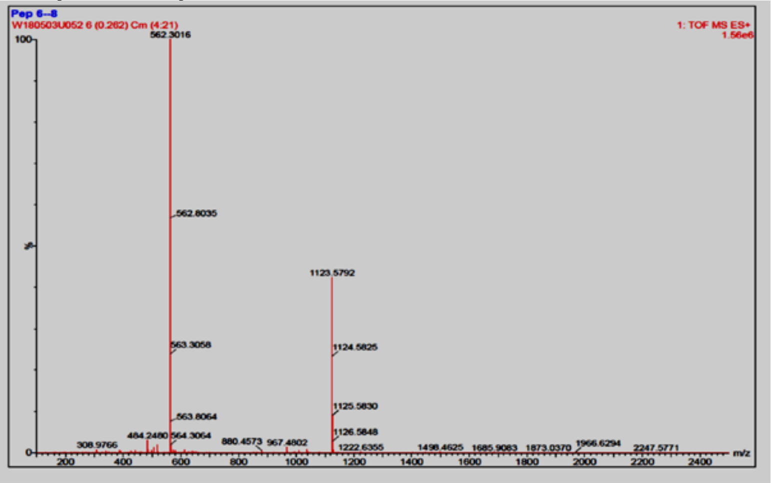


Low-resolution mass spectrum of peptide sythesized gp-100

**Peptide MART-1 Sequence:** ELAGIGILTV
**Chemical Formula:** $C_{45}H_{80}N_{10}O_{14}$
**Molecular Weight:** 985.18

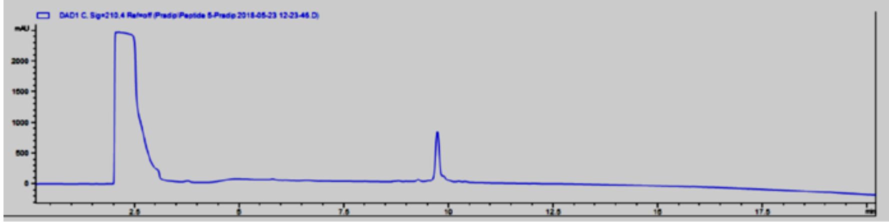

Analytical HPLC profile peptide MART-1. Retention time = 9.80 min (monitored at 210 nm). Purity > 90% by HPLC

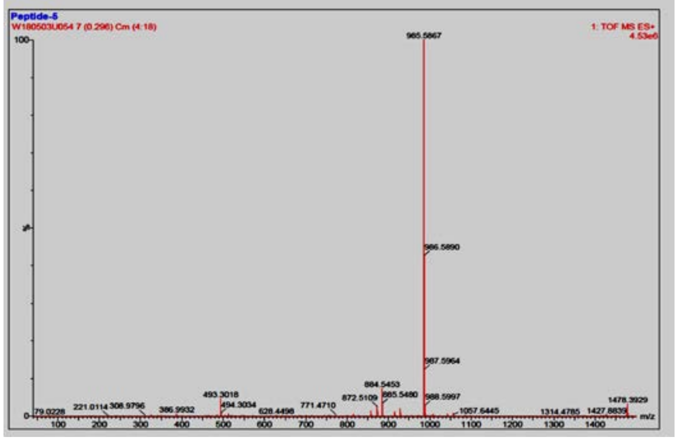


Low-resolution mass spectrum of peptide MART-1
